# Targeting the dysfunctional mitochondrial respiration in drug resistant B-cell acute lymphoblastic leukemia

**DOI:** 10.1101/595959

**Authors:** Rajesh R. Nair, Debbie Piktel, Patrick Thomas, Quincy A. Hathaway, Stephanie L. Rellick, Pushkar Saralkar, Karen H. Martin, Werner J. Geldenhuys, John M. Hollander, Laura F. Gibson

## Abstract

Reprogramming of cellular pathways is a crucial mechanism of drug resistance and survival in refractory acute lymphoblastic leukemia (ALL) cells. In the present study, we performed an unbiased gene expression analysis and identified a dysfunctional mitochondrial respiration program in drug-resistant ALL cells grown in a co-culture system with bone marrow stromal cells (BMSC). Specifically, the activity of the complexes within the electron transport chain was significantly downregulated, correlated with decreased mitochondrial mass and ATP production in drug-resistant ALL cells. To validate mitochondrial respiration as a druggable target, we utilized pyrvinium pamoate (PP), a known inhibitor of mitochondrial respiration and documented its anti-leukemic activity in several ALL cell lines grown alone or in co-culture with BMSC. To increase the bioavailability profile of PP, we successfully encapsulated PP in a nanoparticle drug delivery system and demonstrated that it retained its anti-leukemic activity in a hemosphere assay. PP anti-leukemic activity was decreased by the addition of sodium pyruvate, and furthermore, PP was found to have an additive anti-leukemic effect when used in combination with rotenone, a mitochondrial complex I inhibitor with activity similar to PP on the mitochondrial respiration. Importantly, PP’s cell death activity was found to be specific for leukemic cells as primary normal immune cells were resistant to PP-mediated cell death. In conclusion, we have demonstrated that PP is a novel therapeutic lead compound that counteracts the respiratory reprogramming found in refractory ALL cells.

## Introduction

B lineage acute lymphoblastic leukemia (ALL) originates from differentiation arrest followed by uncontrolled proliferation of the immature B lymphoid cells, resulting in accumulation within the bone marrow (BM) [1]. This rapid accumulation of the leukemic blast cells leads to the “hijacking” of the BM niche and the impairment of steady state hematopoiesis. Thus, coincident with progression of disease, there is a collapse of the functional integrity of the hematopoietic system [2]. The drivers of the disease have not yet been fully elucidated, but are known to include changes in chromosomal number (hyperdiploidy, hypodiploidy, trisomy 4 and 10 and intrachromosomal amplification of chromosome 21) or chromosomal translocation (MLL-AF4, ETV6-RUNX1, E2A-PBX1 and BCR-ABL1) [3]. Fortunately, irrespective of the genesis of the disease, there has been continuous improvement in prognosis in both children and adults, with most achieving complete remission following treatment using the present standard-of-care [4]. However, disease relapse does occur in specific patients, and often the emerging disease will manifest a drug resistant phenotype, thus drastically reducing the chances of complete disease remission and cure in these patients [5, 6]. One of the single most important prognostic indicators for refractory/relapsed disease is the presence of minimal residual disease (MRD) within the BM after completion of the therapeutic regimen [7, 8]. Naturally, identification of therapeutic regimens that specifically target the refractory cells would lead to better therapeutic outcomes for patients who have undergone remission followed by relapse due to incomplete eradication of the MRD from the bone marrow.

In order to delineate the characteristics of the leukemic cells that constitute MRD, we previously developed a co-culture model using primary human BM stromal cells (BMSC) or osteoblast cells with ALL cells [9]. The ALL cells formed three populations including the cells floating in suspension (S) above the stromal cell layer, the cells loosely adhered to the top of the stromal cells and the cells that bury underneath the stromal cells, which appear phase dim (PD) by phase contrast microscopy. Specifically, we demonstrated that the PD cells within the co-culture were more refractory to the present standard-of-care therapies when compared to S cells within the same co-culture [10]. Interestingly, the drug resistant phenotype in PD cells was accompanied by a reduced proliferation index and increased expression of p27 (CDKN1B) resulting in the accumulation of these cells in the G0/G1 phase of cell cycle. Furthermore, PD cells had an altered metabolic profile with increased expression of hexokinase 1 and 2, correlated with heightened glycolytic activity [10]. Additional studies focused on identifying the mechanisms by which BM microenvironment-derived cues modulate resistance in PD tumor cells demonstrated that down-regulation of miR-221/222 and BCL6 were two contributing factors important for quiescence and cell survival in the BM [11, 12]. These studies highlighted the importance of understanding the mechanisms involved in the BM microenvironment-mediated protection of leukemic cells leading to MRD in ALL. More importantly, the results demonstrated the utility of studying the drug resistant PD cells as a model to interrogate novel therapeutic strategies that could effectively eradicate refractory cells within the BM.

In the present study, we performed unbiased gene expression profiling of the leukemic PD cells in co-culture with BMSC, and observed that their mitochondrial function was altered compared to leukemic cells growing alone in media (M). We have further characterized this functional alteration and hypothesized that mitochondrial bioenergetics offers an actionable target for anti-leukemic therapy. In this study, we used pyrvinium pamoate (PP) which targets mitochondrial respiration [13], to test this hypothesis. The choice of PP is based on its demonstrated anti-leukemic activity in nine different human-derived ALL cell lines of varied phenotypes. Furthermore, this anti-leukemic activity was, in part, associated with targeting and disruption of leukemic mitochondrial respiration. Based on these data, we suggest that agents like PP that target mitochondrial respiration show promise as anti-leukemic therapies. As a novel lead compound, PP, may be useful in combination with the present standard-of-care to treat relapsed and refractory ALL.

## Experimental procedures

### Materials and cell culture

TOM1 (DSMZ ACC#578), SUPB15 (ATCC #CRL-1929), and JM1 (ATCC #CRL-10423) were purchased and maintained in RPMI 1640 supplemented with 10 % FBS, 0.05 mM β-mercaptoethanol and 1x streptomycin/penicillin antibiotics. REH (ATCC #CRL-8286), NALM1 (ATCC #CRL-1567), NALM6 (DSMZ ACC #128), BV173 (DSMZ ACC#20), RS4 (ATCC #CRL-1873) and SD1 (DSMZ ACC#366) were purchased and maintained in RPMI 1640 supplemented with 10 % FBS and 1x streptomycin/penicillin antibiotics. De-identified primary BMSC were provided by the WVU Cancer Institute Biospecimen Processing Core and the WVU Department of Pathology Tissue Bank. ALL cell lines were authenticated by short tandem repeat (STR) analysis (University of Arizona Genetics Core, Tucson, AZ) and maintained in 6 % CO_2_ in normoxia at 37°C. Pyrvinium pamoate (PP) was purchased from Sigma Aldrich (Cat #P-0027) and stored at −80°C as a 10 mM stock.

### Long-term co-culture and isolation of leukemic cell population

Long-term co-culture conditions were followed as previously described [9]. Briefly, 1 million ALL cells were seeded on an 85% confluent BMSC layer and maintained in 5% O_2_. The co-culture was fed every 4 days and PP was added, where indicated, on the 8th day of culture. On the 12th day in culture, the ALL cells were isolated for further processing. The leukemic cell population that was in suspension and not interacting with the stromal cells was collected and designated as suspended cells (S). The leukemic cells which were buried under the BMSC were separated by size exclusion with G10 sephadex after vigorous washing to remove all leukemic cells adhered to the top of the BMSC [9]. Buried leukemic cells were designated phase dim cells (PD) and have been previously described to be the most chemotherapy-resistant population [11, 12]. As such, they are not assumed to be identical, but rather are used as a model, for refractory tumor cells that are known to be clinically problematic in the treatment of ALL.

### RNA sequencing

RNA was isolated using a Qiagen RNAeasy mini kit and sent to the West Virginia University Genomics Core for next-generation sequencing. Libraries were prepared using polyadenylation selection with the KAPA stranded mRNA-Seq kit. Samples were sequenced as paired-end reads across two lanes using the Illumina HiSeq. Fastq files were first trimmed of any 3’ adapters using Cutadapt and then aligned to the human reference genome GRCH38/hg38 with HISAT2 [14, 15]. Rsubread was implemented for read counting [16]. After read counting, all analyses were conducted within the R environment. Sample metrics were evaluated using both NOISeq [17, 18] and DESeq2 [19], though only DESeq2 was employed for differential expression analysis. The data has been deposited: SRA accession: PRJNA509768.

### Electron transport chain complex activities

The activity of the ETC Complexes I, III, IV and V were measured as previously described [20]. Briefly, SUPB15 or REH cells in media, or suspended in co-culture (S) or buried in co-culture (PD), were processed and treated with RIPA buffer. The reduction of decylubiquinone in the lysate was used as a measure of Complex I activity, reduction of cytochrome c was used as a measure of Complex III, oxidation of reduced cytochrome c was used as a measure of Complex IV and finally ATP synthase activity was measured as oligomycin-sensitive ATPase activity through pyruvate kinase and phosphoenolpyruvate. Protein quantification was performed to normalize the activity measurement of each complex.

### ATP measurement

A luminescent ATP detection assay kit was used to determine the total levels of cellular ATP as per manufacturer’s instructions (abcam, Cat #ab113849). Briefly, long-term co-cultures were either treated with 125 nM of PP or left untreated followed by isolation of the different cell populations, as described above. 20,000 of the isolated cells were then lysed and added into the detection solution and the luminescence was measured using a BioTek plate reader.

### Mitochondrial staining assay

Leukemic cell populations were isolated from long-term co-culture as described above. The isolated cells were then incubated with 100 nM of MitoTracker Deep Red FM (Invitrogen, Cat# M22426) for 30 min. After a single wash with PBS, the cells were fixed with 3.7% paraformaldehyde and the MitoTracker intensity was ascertained using flow cytometry (BD LSR Fortessa).

### Antibodies and western blot analysis

Monoclonal anti-β-Catenin antibody was purchased from Cell Signaling Technology (Cat #8480) and used at 1:1,000 dilution. Mouse polyclonal anti-GAPDH was purchased from Research Diagnostics Inc and used at 1: 20,000 dilution. Cells were lysed using RIPA buffer and the resultant protein concentrations were determined using the bicinchoninic acid (BCA) protein assay kit (Pierce, Cat #23227). Equal quantities of proteins were resolved on SDS-PAGE gels and transferred to nitrocellulose membranes. Membranes were blocked in TBS with 5 % nonfat dry milk and 0.5 % Tween-20 and probed with the indicated primary antibodies. After incubation with horseradish peroxidase-conjugated secondary antibodies, signal was visualized using Amersham ECL Prime western blot detection reagent (GE Life technologies, Cat # RPN2232). Western blots are representative of at least two independent experiments.

### Cell proliferation assay

Cells were plated in 96-well clear plates at a density of 50,000 cells per well. Cells were treated with PP at indicated concentrations and the cell proliferation was tested after 72 hrs of incubation with the compounds. A cell counting kit was utilized according to the manufacturer’s instructions (Dojindo Molecular Technologies Inc., Cat # CK04). Briefly, 10 μl of the assay reagent was added to each well and incubated for 2 hrs at 37°C, after which the plates were read on a BioTek Cynergy 5 plate reader at 450 nm absorbance. Untreated cells were used as controls.

### Nanoparticle encapsulation of PP

The nanoparticles were prepared as described earlier [21]. Briefly, 2 mg of PP was dissolved in 500 µl of acetone, along with 15 mg PEG-PLGA-COOH, and added dropwise into a 0.5 % w/v solution of polyvinyl alcohol (PVA). The primary emulsion was formed by probe sonication (Heat Systems Ultrasonics Inc., Model W-225) for 10 minutes at an output level of 6 using a pulsed mode. The nanoparticles were obtained after evaporating acetone by stirring, over a period of 24 hrs at room temperature. The particle size of the nanoparticles was analyzed using a Nanosight NS300 (Malvern Panalytical, Malvern, UK). The nanoparticle samples were diluted prior to the measurements. The zeta potential of the blank and drug loaded nanoparticles was measured using the Malvern Zetasizer (Malvern, UK) (Fig. S6).

### Cell death analysis

Cells were grown in 24-well plates at a density of 250,000 cells per well. Cells were treated with 125 nM of PP or 1 μM of rotenone for 24 hrs. Cells were stained using Annexin V-FITC according to manufacturer’s instructions (ThermoFisher Scientific, Cat# 88-8005) and analyzed using flow cytometry (BD LSR Fortessa).

### Hemosphere assay

SD1 cells were plated at 100,000 cells/well in a 96 well plate and allowed to form spheroids for 4 days. At the end of 4 days the formation of spheroids was visualized and confirmed using light microscopy. The spheroids were then treated with 250 nM of nanoparticle encapsulated PP, and the resulting effects on the spheroids were captured using a Leica camera attached to a Leica DMIL LED microscope.

### Statistical analysis

All the experiments were carried out as three independent experiments unless specifically noted otherwise in the legends. All the data were represented as mean ± SEM, and p<0.05 was considered statistically significant. Statistical significance between two groups were carried out by using student’s T-test. For comparing three or more groups, a one-way ANOVA was used to determine statistical significance followed by a post-hoc Tukey’s test. For the gene expression analysis data, DESeq2 was used to calculate significance through implementation of the Wald test. In brief, adjusted p-values are derived from both the unadjusted p-value and False Discovery Rae (FDR). DESeq2 uses a generalized linear model of computing significance across each sample, where a negative binomial model is fitted to the counts. The FDR in DESeq2 follows the Benamini-Hochberg procedure.

## Results

### Phase dim (PD) leukemic cells in co-culture have dysfunctional mitochondrial respiration

To characterize the resistant phenotype of the refractory ALL cells, we used our long-term co-culture model of REH cells as a representative ALL cell line with BMSC to isolate the transcribed RNA from the phase dim (PD) population for comparison to transcribed RNA from cells in suspension (S) or in media alone (M). Sample-to-sample distribution of transcript counts (Fig. S1A and B) illustrate a distinctive phenotypic shift in the REH PD cells following treatment, attributable to changes in global genomic expression and not to RNA content (Fig. S1C). As shown in Fig. S1D, compared to the M cells, the PD cells differentially expressed roughly 2,600 genes. Interestingly, no difference in gene expression profile was observed between the PD cells and S cells (Fig. S2A). Furthermore, hexokinase II (HK2) was found to be the most significantly over-expressed in the PD cell population in all the four independent co-cultures that were used for the gene expression analysis (Fig. S2B), which validated the cell culture conditions and the resistant phenotype of the REH PD cells. Since HK2 is an important player in mitochondrial respiration, we utilized the Ingenuity Pathway Analysis (IPA) software to explore mitochondrial respiratory pathways that were differentially regulated. The oxidative phosphorylation module was the most significantly down-regulated pathway in the PD cells (Fig. 1A). Also, looking specifically at the mitochondrial oxidative phosphorylation pathway revealed that expression of genes in Complex I, Complex III, Complex IV and Complex V of the electron transport chain (ETC) were down-regulated in the PD cells, while Complex II remained transcriptionally unaffected (Fig. 1B).

**Figure 1.**
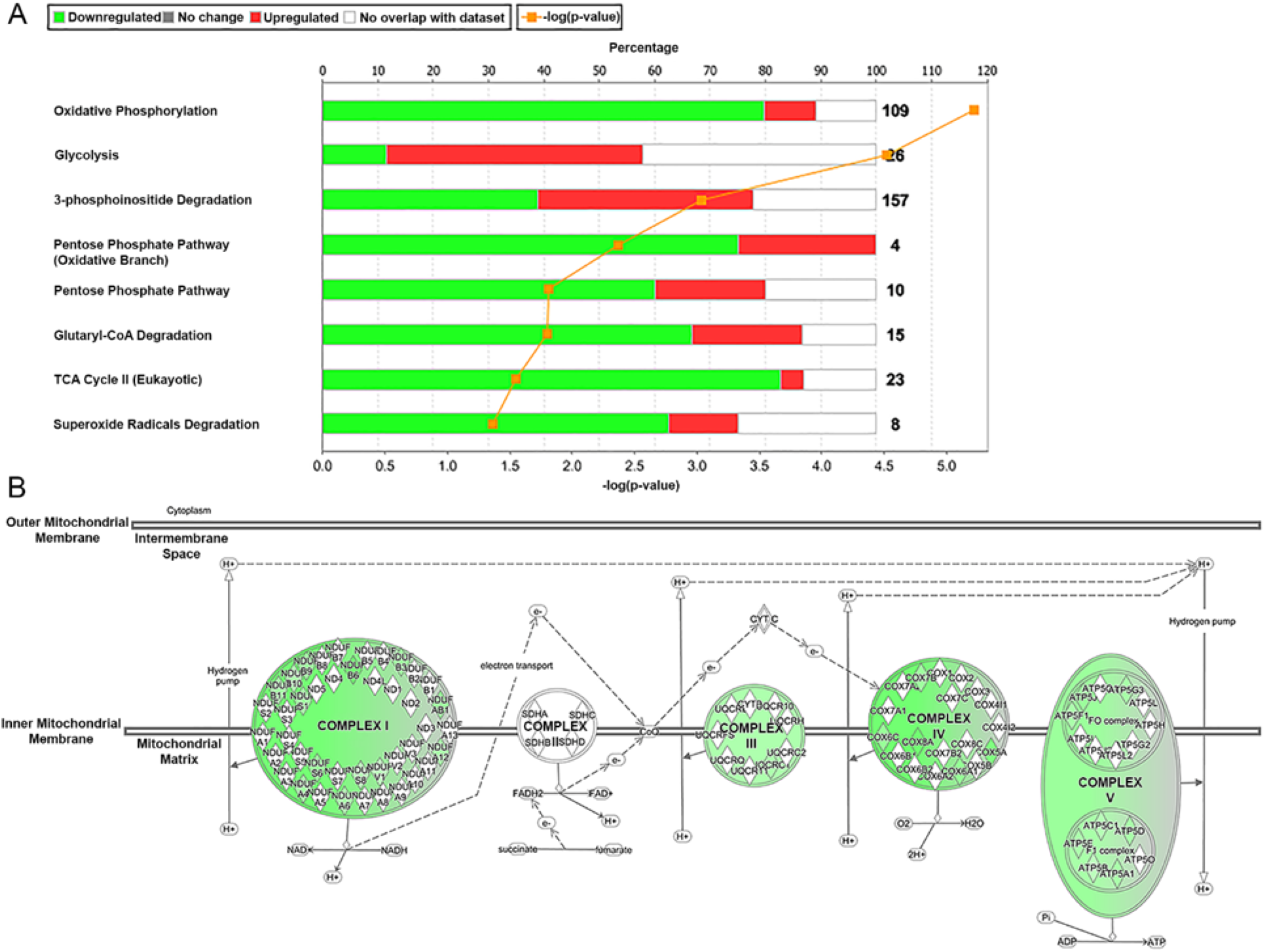
Mitochondrial deregulation in REH PD cells in co-culture with BMSC. (A) The top eight altered mitochondrial pathways, determined by cumulative p-values within a pathway along with the size of the pathway (orange line). The number next to each bar indicates the proteins in each pathway, while the colored legends describes the changes in expression. (B) IPA pathway examining changes to mitochondrial complex transcripts. Green indicates significantly decreased expression in the PD cells compared to cells grown in media. Grey indicates no change in gene expression.

### Co-culture affects activity of mitochondrial complexes

To validate the observations derived from the gene expression analysis, we assayed the enzymatic activity of the mitochondrial complexes in two ALL cell lines (REH and SUPB15) using cell lysates from M, S and PD cells. Complex I activity of the S and PD cells was significantly reduced when compared to M cells for the REH ALL cell line; while it was low but unchanged in all three populations of the SUPB15 cells (Fig. 2A). Specifically, Complex I activity was decreased by 58 % in S cells and by 55 % in the PD cells as compared to M cells. Interestingly, for SUPB15 cells, the PD cells showed significant reduction in Complex III activity compared to M and S cells (Fig. 2B). Furthermore, Complex IV and Complex V showed no significant differences in activity between the three cell populations in either REH or SUPB15 cells (Fig. 2C and D).

**Figure 2.**
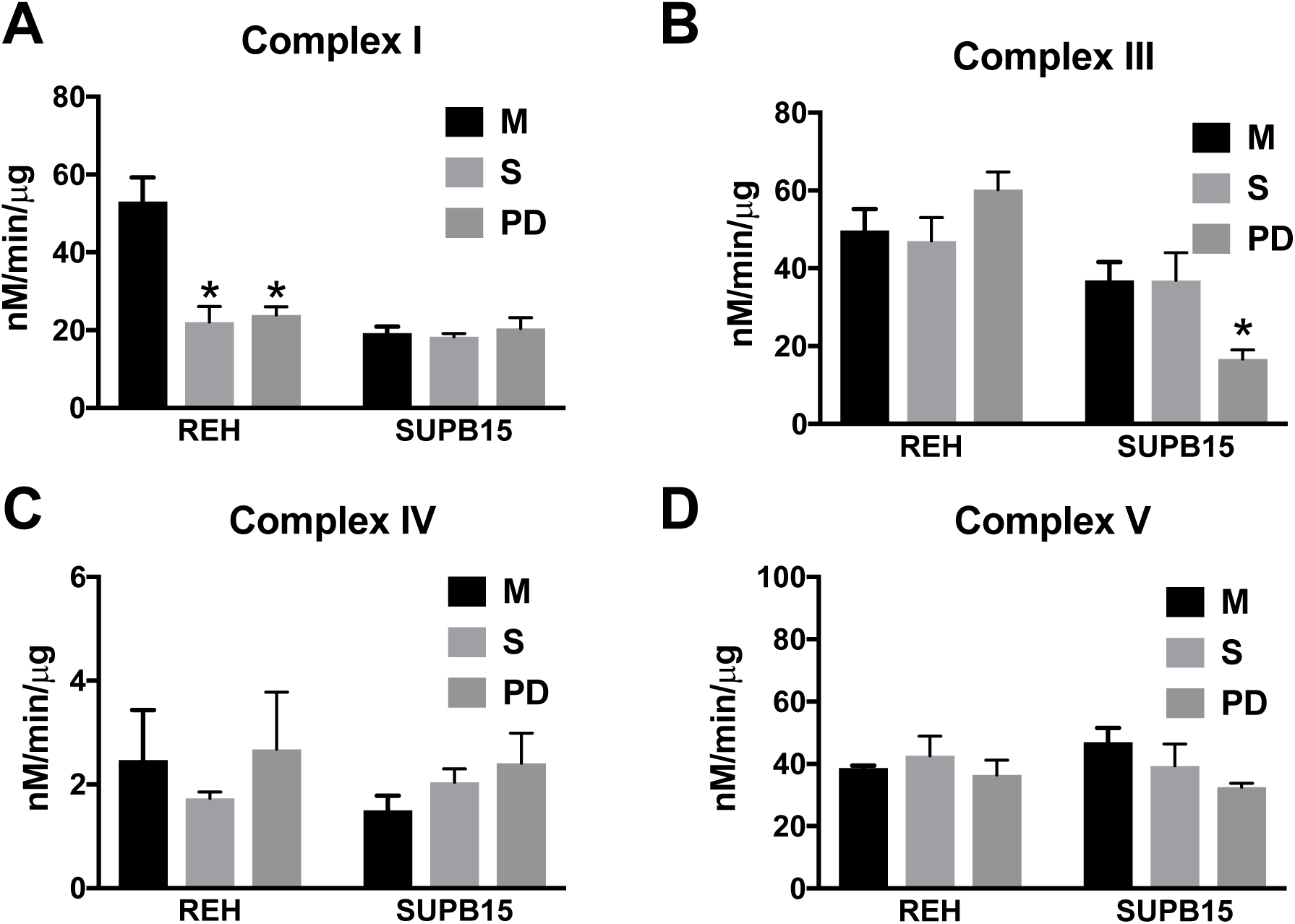
Electron transport chain complex activity in leukemic cells in co-culture with BMSC. REH or SUPB15 cells were maintained in long-term co-culture with BMSC as described in the methods. Following 12 days in culture, leukemic cells growing in media alone (M), suspended leukemic cells from co-culture (S) and phase dim leukemic cells from co-culture (PD) were collected and the activities of Complex I (A), Complex III (B), Complex IV (C) and Complex V (D) were measured. The data are presented as mean ± SEM and represent three experiments conducted independently. * p<0.05, as compared to cells grown in media (M).

### Leukemic cells in co-culture have dysfunctional mitochondrial bioenergetics

To understand how the decreased ETC complex activity of the leukemic cells in co-culture affects the mitochondrial bioenergetics, we looked at ATP production and the relative mitochondrial mass within the different sub-populations of leukemic cells in co-culture. We found that the ATP production was significantly decreased in both REH and SUPB15 cells in co-culture with BMSC when compared to the M cells (Fig. 3A and B). Interestingly, while the SUPB15 sub-population of S and PD cells showed similar (∼18 %) decrease in ATP production, in REH cells the PD cells had a greater decrease in ATP production (∼42 %) than the S cells (∼19 %) when compared to the REH M cells. Furthermore, MitoTracker fluorescence intensity in the resistant PD cells was significantly lower when compared to the S or M cells (Fig. 3C and D). This decrease in mitochondrial staining was very specific and similar in PD cells from both the tested cell lines (30 % in REH PD cells and 40 % in SUPB15 PD cells compared to their respective cells in media).

**Figure 3.**
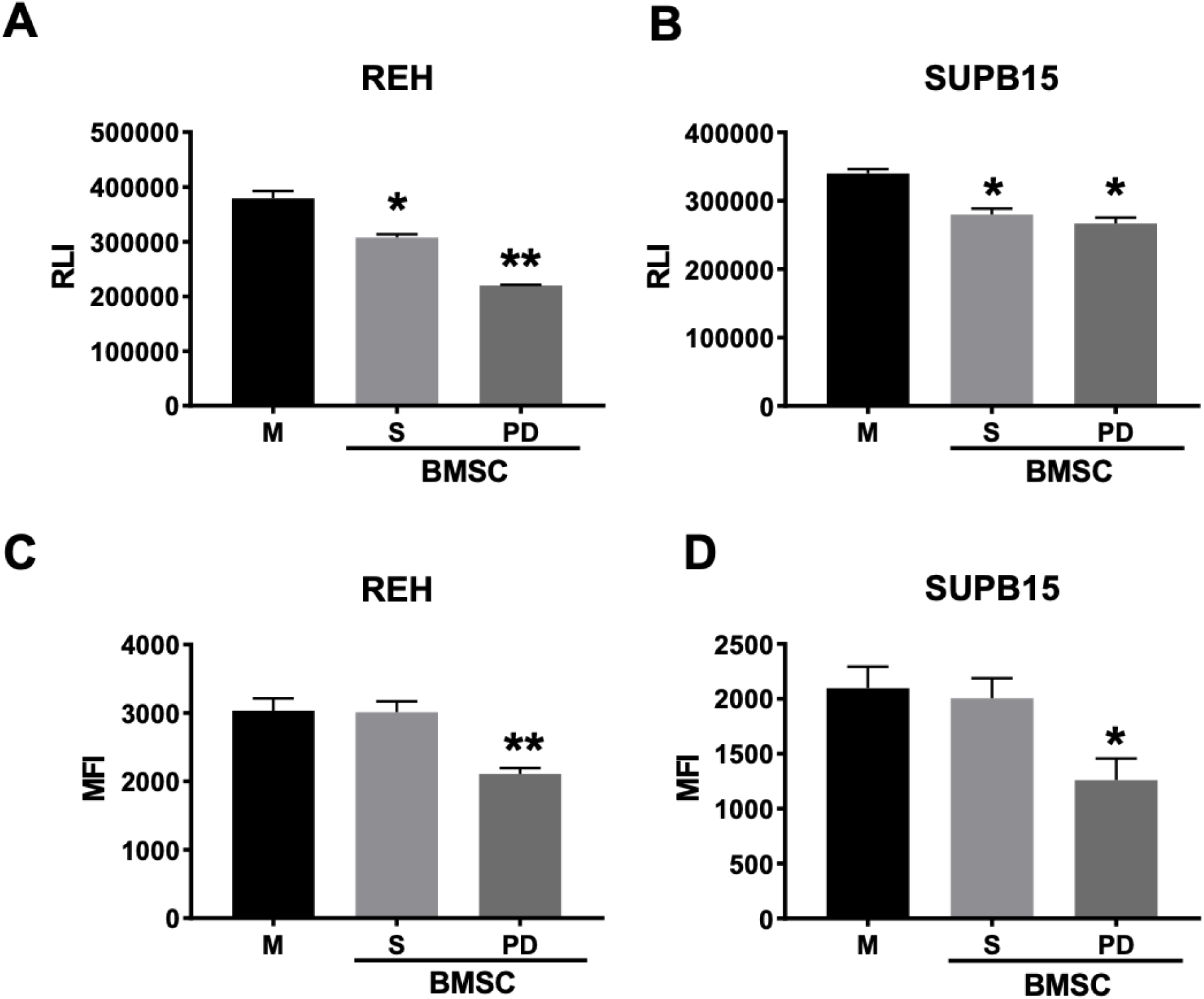
Characterization of mitochondrial function and mass in ALL cells in co-culture with BMSC. REH (A & C) and SUPB15 (B & D) were maintained in long-term co-culture with BMSC as described in the methods. Following 12 days in culture the ALL cells growing alone (M), suspended cells (S) and the phase dim cells (PD) were isolated and either subjected to ATP detection and quantitated as relative luminescence intensity (RLI) (A & B) or mitochondrial staining and quantitated as mean fluorescence intensity (MFI) (C & D) as described in the method section. The data is presented as mean ± SEM and is a representative of a single experiment done in triplicate and performed three independent times. * p<0.05, when compared to cells grown in media (M), ** p<0.05, when compared to cells grown in media (M) or the cells in suspension in co-culture (S).

### Pyrvinium pamoate (PP) demonstrates anti-leukemic effects in ALL cells

Since our previous results showed that the mitochondrial respiration was severely downregulated in the PD cell subpopulation, we aimed to determine if it could serve as a therapeutic target. We used PP to inhibit mitochondrial respiration in a panel of ALL cell lines (15). Treatment with PP decreased cell proliferation in a concentration-dependent manner in all eight of the cell lines cultured in media without microenvironment cues (BV173, JM1, NALM1, NALM6, REH, RS4, SUPB15 and TOM1) (Fig. 4A). The IC50 of PP ranged from 0.17 μM for REH cells to 1 μM for RS4 cells (Fig. 4B). To determine the effect of PP on the cell cycle, we treated REH and SUPB15 cells with PP and observed a significant decrease in the number of cells in the S and M phases of the cell cycle (Fig. S3A and B). Subsequently, we wanted to evaluate the effects of PP on leukemic cells when they were co-cultured with BMSC. Treatment with PP resulted in a decrease in the total number of live cells in both REH and TOM1 cell lines, irrespective of cell culture conditions (Fig. 4C and E). This overall decrease in the total number of live cells was accompanied by a significant decrease in cell viability. Specifically, in REH cells, a ∼55% decreased viability in M and S cell populations was observed compared to ∼80 % in the PD cell population (Fig. 4D). A similar pattern was observed in TOM1 cells treated with PP, with a ∼75% decreased viability in M and S cell population compared to ∼97% viability in the PD cell population (Fig. 4F).

**Figure 4.**
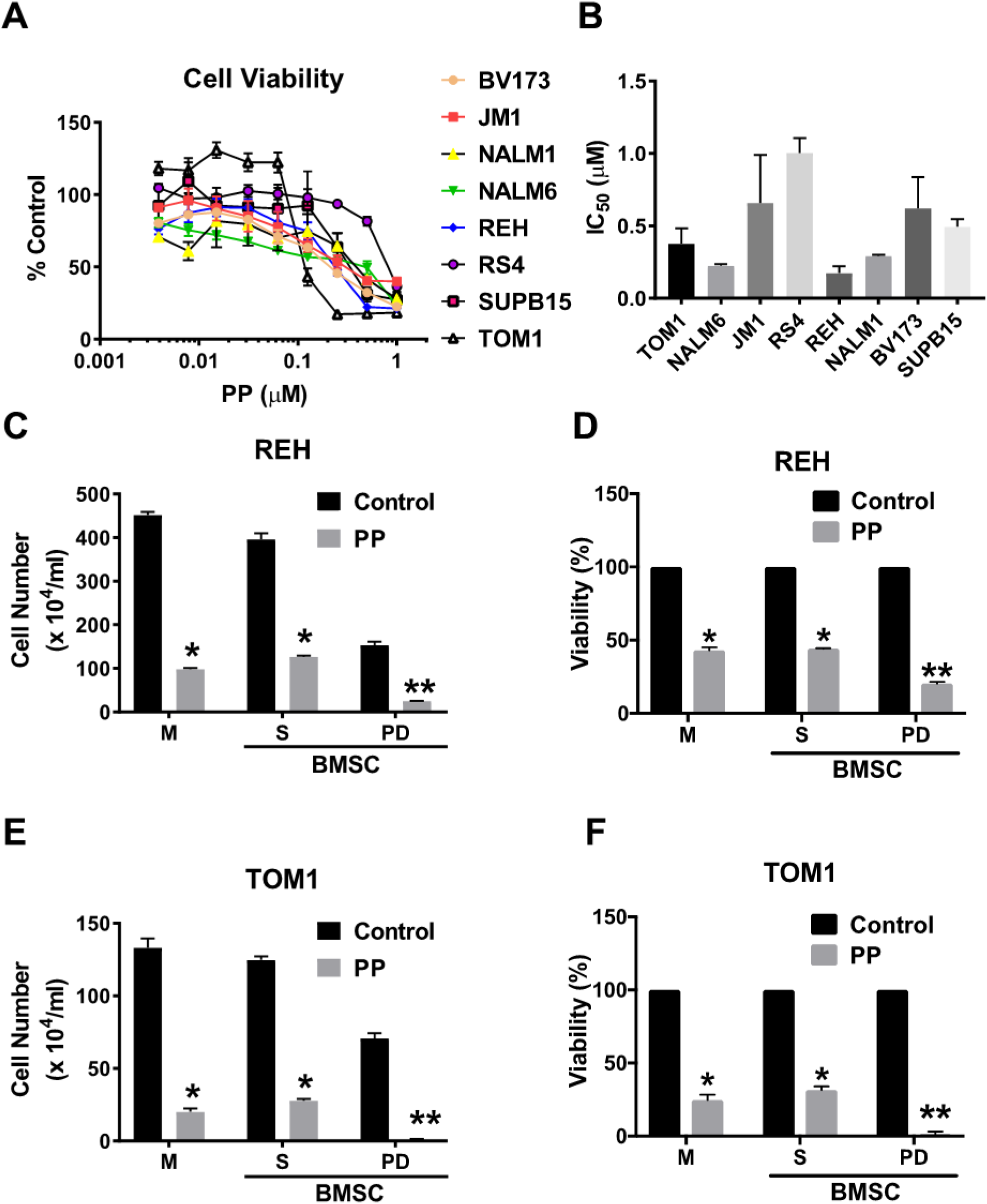
Anti-leukemic activity of pyrvinium pamoate. (A) Indicated ALL cell lines were plated at 50,000 cells/well in a 96 well plate and treated with increasing concentrations of pyrvinium pamoate for 72 hrs. Cell proliferation was measured using a cell counting kit as described in the methods. (B) IC_50_ values were calculated from the dose response curve for each cell line using the compusyn software. REH (C & D) and TOM1 (E & F) were grown in co-culture with BMSC or alone (M) in a 24 well plate for 12 days. On day 9 in co-culture, the cells were treated with 250 nM pyrvinium pamoate (PP). The live cells were counted using Trypan blue dye exclusion method for cells grown alone (M), cells in suspension (S) and cells buried (PD) when in co-culture in REH (C) and TOM1 (E). The total cells and the live cell population was used to compute the % viability in REH (D) and TOM1 (F). The data is presented as mean ± SEM and is a representative of a single experiment done in triplicate and performed three independent times. * p<0.05, when compared to corresponding untreated group, ** p<0.05, when compared to all other untreated and treatment groups.

### Pyrvinium pamoate nanoparticles (NP [PP]) demonstrate anti-leukemic activity

Since PP has poor bioavailability, we encapsulated it within nanoparticles composed of PGLA and PEG complexes. We first determined if the nanoparticles (NP) by themselves had any activity on the bone marrow stromal components or the ALL cells. As shown in Fig. 5A, NP at doses as high as 100 μM failed to elicit any death in BMSC, osteoblasts (HOB), NALM27 or REH ALL cells lines. Next, we tested the toxicity of the PP encapsulated in NP in normal cells compared to ALL cells. Fig. 5B shows that 250 nM of the PP encapsulated in NP did not have any deleterious effects on BMSC, normal bone marrow mononuclear cells (BMMC), or normal peripheral blood mononuclear cells (PBMC). However, it did cause a significant decrease in cell proliferation (46 %) when exposed to a B lymphoblastoid cell, SD1 (Fig. 5B). To ascertain that the differential effects of PP between the normal and the leukemic cells was not due to differences in cell proliferation rate, we utilized normal primary CD3+ cells and induced proliferation using a CD3/CD28 T cell activator. Treatment of dormant or proliferating T cells with PP did not result in decreased cell viability in the dormant T cell population and showed a minimal decrease in viability (∼13 %) in the proliferating T cells (Fig. S4). Finally, we wanted to determine the anti-leukemic effects of the PP encapsulated NP in a hemosphere assay. We utilized SD1 cells to form hemospheres for 4 days, after which they were treated with 250 nM of the PP encapsulated NP. After 48 hrs of treatment, the NP were able to penetrate the pre-formed spheres resulting in a loss of hemospheres (Fig. 5C). Quantitation of the spheres using light microscopy showed that the PP encapsulated nanoparticles resulted in a loss of spheres and were efficient in inhibiting the sphere-forming abilities of SD1 cells, to the point no spheres formed after treatment (Fig. 5C).

**Figure 5.**
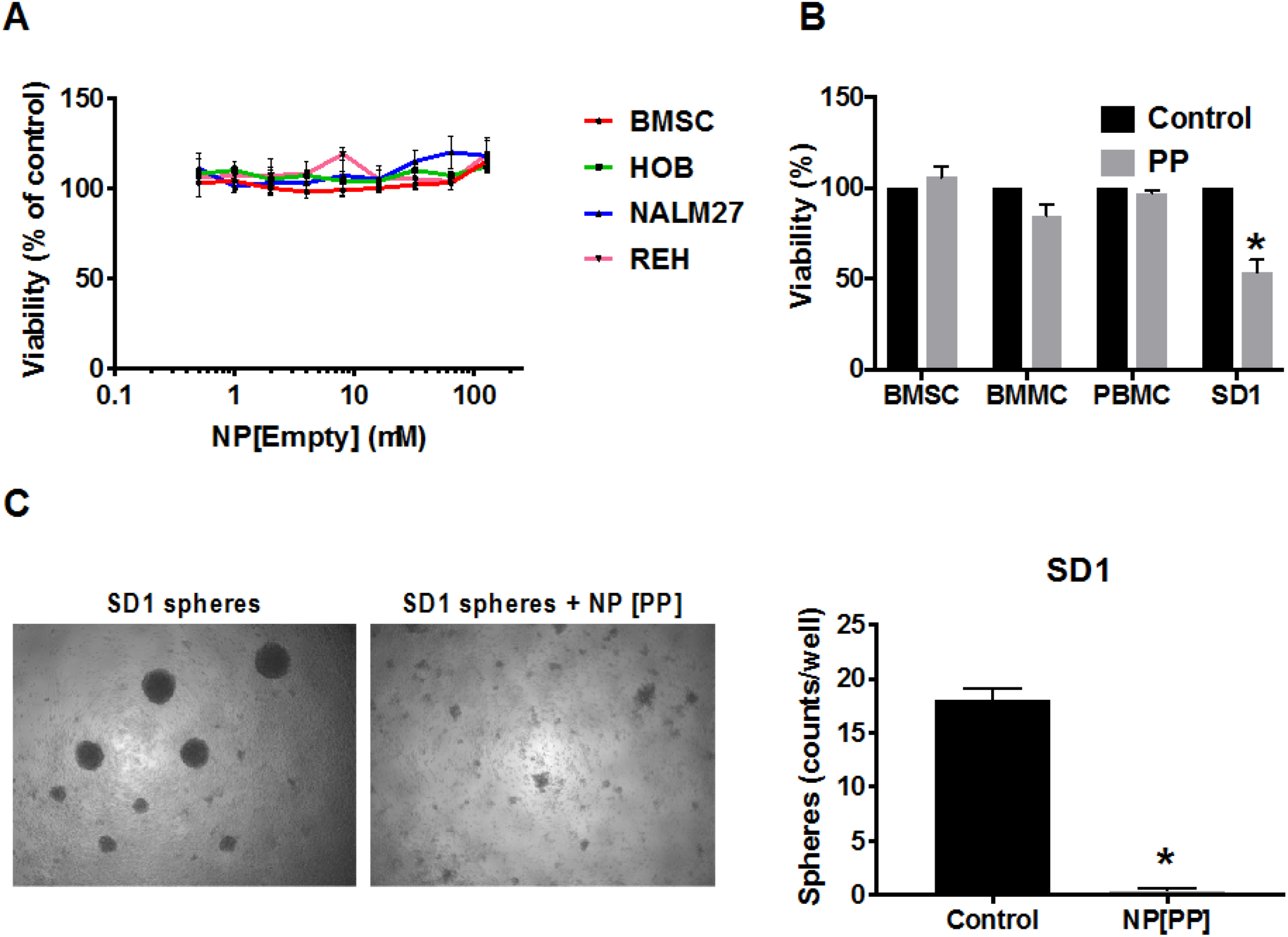
Pyrvinium pamoate encapsulated in nanoparticle demonstrates anti-leukemic activity. (A) BMSC, normal human osteoblasts (HOB), and ALL cell lines NALM27 and REH were plated at 50,000 cells/well in a 96 well plate and treated with increasing concentrations of empty nanoparticles (NP[Empty]) for 72 hrs. Cell proliferation was measured using a cell counting kit as described in the methods. (B) BMSC, normal bone marrow mononuclear cells (BMMC), normal peripheral blood mononuclear cells (PBMC), and an ALL cell line (SD1) were plated at 50,000 cells/well in a 96 well plate and treated with 250 nM pyrvinium pamoate encapsulated in nanoparticles for 72 hrs. Cell proliferation was measured using a cell counting kit as described in the methods. (C) Light microscope images of SD1 hemospheres that were allowed to form for 4 days and then treated with 250 nM of pyrvinium pamoate encapsulated in nanoparticles (SD1 sphere + NP [PP]) for 48 hrs. Untreated spheres were used as control. The data is presented as mean ± SEM and is a representative of a single experiment done in triplicate and performed three independent times. * p<0.05, when compared to all other untreated and treatment groups.

### Pyrvinium pamoate encapsulated nanoparticles inhibit mitochondrial respiration

To understand the mechanism of the PP-induced cell death in ALL cells, we analyzed its effect on ATP production. As seen in Fig. 6A & B, treatment with 250 nM PP for 24 hrs significantly reduced the ATP production in both REH (70 %) and SUPB15 (93 %) cell lines compared to controls. Sodium pyruvate has been shown to facilitate glycolysis in cells that have dysfunctional mitochondria [22]. To determine if the PP-induced death in leukemic cells could be diminished by addition of sodium pyruvate, we grew the leukemic cells in glucose-free media and supplemented the media with 10 mM sodium pyruvate. As seen in Fig. 6C & D, compared to PP-treated cells growing in glucose-free media, PP-mediated cell death was significantly decreased in the cells supplemented with sodium pyruvate. Further, to ascertain whether PP-induced death in ALL cells was solely dependent on inhibition of mitochondrial respiration, we pretreated REH and SupB15 with 1 μM rotenone, an inhibitor of Complex I of the ETC [23]. The pretreatment of rotenone was followed by exposure of cells to 150 nM of PP. As seen in Fig. 6E, pretreatment of rotenone caused 20% cell death in REH cells and treatment with PP encapsulated NP caused 60% cell death in REH and both given together showed an additive 80% cell death in REH cells. Similarly, in SUPB15 cells, rotenone induced 20% cell death and PP encapsulated NP caused 20 % cell death in the same cells. Finally, when given together, PP and rotenone induced an additive 40% cell death (Fig. 6F).

**Figure 6.**
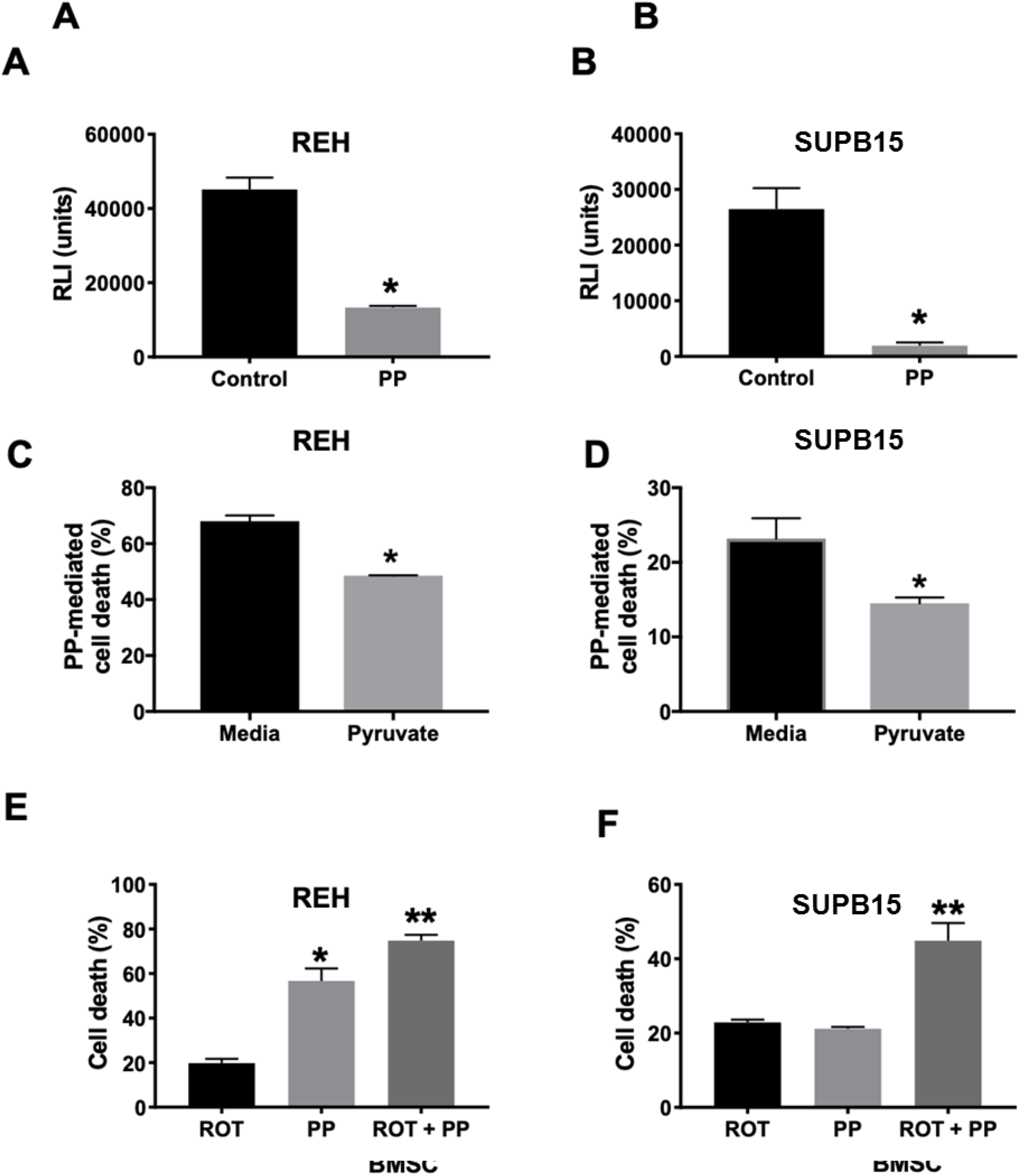
Pyrvinium pamoate’s anti-leukemic activity requires inhibition of mitochondrial respiration. REH cells (A) or SUPB15 cells (B) were plated at 250,000 cells/well in a 24 well plate and then treated with 250 nM pyrvinium pamoate (PP) for 24 hrs. Following treatment, the total ATP was measured and quantitated as the relative luminescence intensity (RLI) as described in the methods. REH cells (C) or SUPB15 (D) cells were grown overnight in glucose-free media or glucose-free media supplemented with 10 mM sodium pyruvate. After overnight incubation, the cells were treated with 250 nM of PP and cell death was analyzed after 24 hrs using flow cytometry. REH cells (E) or SUPB15 (F) were pretreated for 2 hrs with rotenone and followed by treatment with pyrvinium pamoate (ROT + PP) for 24 hrs. Cells treated with rotenone alone (ROT) or pyrvinium pamoate alone (PP) were used for comparison. Untreated cells were used as control to measure drug-mediated cell death by annexin V binding assay. The data is presented as mean ± SEM and is a representative of a single experiment done in triplicates and performed three independent times. * p<0.05, when compared to all other untreated and treatment groups.

## Discussion

In the present study, we have utilized an in vitro co-culture system to characterize PD drug-resistant leukemic cells that mimic the cell population that constitutes minimal residual disease found within the BM of relapsed ALL patients. We have shown that the gene expression profile of proteins involved in the mitochondrial ETC complex is downregulated in PD cells, with a significant reduction in Complex I activity, and that mitochondrial dysfunction was accompanied by a reduction in mitochondrial mass and reduced ATP production in PD cells. PP inhibits mitochondrial bioenergetics and induces cell death in a variety of ALL cell lines and importantly is also able to specifically target the resistant PD cells for death. Looking ahead toward possible use in the clinic, we also showed that PP encapsulation in nanoparticles to potentially increase its bioavailability does not diminish the activity of PP and opens up an avenue to increase its systemic availability. The complete mechanism of PP-mediated leukemic cell death includes its ability to inhibit ATP production in addition to its mitochondrial inhibitory activity. Taken together, this report demonstrates the potential for the use of PP as an anti-leukemic agent, especially in treatment of patients with relapsed refractory disease.

We had previously reported that the drug resistant ALL cell sub-population designated as the PD cells have increased glycolytic programing [10]. This finding has been further validated by studies showing that B-lymphoid transcription factors (PAX5 and IKZF1) that are critical in normal B-cell development are metabolic gatekeepers that negatively regulate energy metabolism and are mutated or deleted in majority of ALL patients [24, 25]. Furthermore, tumors from B cell ALL patients, but not T cell-ALL, have fewer mitochondria than from patients with myeloid lineage leukemia and abnormality/irregularities in the ALL patient’s mitochondria had a direct correlation to poor treatment prognosis [26]. Also, patient ALL CD34+ cells show upregulation of genes that facilitate glycolysis and downregulation of genes related to the tricarboxylic cycle [27]. In the present study, we show that our drug-resistant PD population has the highest induction in HK2, the master regulator of glycolysis [28]. However, we have shown that this comes at the cost of decreased ability of the drug-resistant ALL cells to utilize oxidative phosphorylation for its energy needs. Specifically, we have demonstrated that the activity of mitochondrial Complex I was severely decreased in ALL cells in co-culture with BMSC, and this was accompanied by decreased ATP production and mitochondrial mass within the drug resistant PD cells.

Sensitization of ALL cells to chemotherapy has been achieved recently by the use of an antibiotic, Tigecycline, that acted through inhibition of mitochondrial respiration [29]. However, Tigecycline has limited activity as a single agent and its anti-leukemic activity in resistant cell lines remains to be tested. In our study, we selected PP as our agent of choice for inhibiting mitochondrial respiration in ALL cells. PP has been shown to have activity in both solid tumors like breast cancer and prostate cancer [30, 31] and hematological malignancies like chronic myeloid leukemia and multiple myeloma [32, 33]; however, its activity in ALL has not yet been reported. PP exerts its cytotoxic activity by inhibiting NADH-fumarate reductase found within Complex II of the mitochondrial ETC system [34]. Additionally, PP was also shown to decrease ATP production and mitochondrial respiration by targeting Complex I and Complex III activity [35]. The cytotoxic activity of PP was found to be exaggerated when cancer cells were grown in hypoglycemic conditions, such as that found within the tumor microenvironment [36]. In the present study, we show that PP demonstrates anti-leukemic activity in different ALL cell lines irrespective of their inherent mechanism of malignancy. Indeed, PP was equally sensitive in ALL cells (REH) driven by ETV6-RUNX1 oncogene as well as in ALL cells (TOM1) driven by BCR-ABL oncogene. Furthermore, we were able to show that this anti-leukemic effect was initiated by a rapid fall in ATP levels, which indicated that PP targeted and inhibited mitochondrial respiration. Most importantly, PP was specifically active in the drug resistant PD cells indicating its potential for inducing death in drug resistant cells found within the relapsed refractory patients.

We had previously shown, using a Seahorse metabolic assay, that rotenone induced a complete inhibition of oxygen consumption rate in ALL cells [10]. In our study, pretreatment of ALL cells with rotenone did not abolish the anti-leukemic activity of PP. In fact, the combination of rotenone and PP gave an additive cell death activity indicating a second mechanism of action for inducing death in ALL cells by PP. Our lab had previously reported the importance of β-catenin signaling in not only maintaining the stem cell profile of leukemic stem cells but also in providing proliferative cues that are critical for survival within the BM microenvironment [37, 38]. At the same time, PP has been shown to activate casein kinase 1α and destabilize β-catenin protein accelerating its degradation and, in the process, attenuating wnt signaling in cancer cells [39]. However, this casein kinase 1α activation by PP to inhibit wnt signaling has been disputed in studies showing that GSK-3β activation was more critical for PP-mediated inhibition of wnt [40]. To delineate if inhibition of β-catenin stability was a mechanism involved in PP-mediated death in leukemic cells, we first showed that β-catenin expression was increased in the resistant PD cell population of ALL cell lines (Fig. S5). Importantly, PP was successfully able to reduce the levels of this protein in the PD cells population, indicating that in addition to mitochondrial respiration, reducing β-catenin levels may play a role in the anti-leukemic effects of PP (Fig. S5).

PP is an FDA-approved drug and is safe for human use in an oral dosage form for treatment of pinworms [13]. However, its systemic bioavailability after oral administration is negligent, making it a very poor choice of agent for use in treating leukemia [41]. A study looking at the *in vivo* effect of PP on androgen receptor has shown that intraperitoneal injections of 1 mg/kg results in a peak plasma levels of 150 nM, which is near the IC50 value of PP used in our present study [42]. Unfortunately, this dose was shown to cause toxicity as seen by a rapid weight loss in the treated mice. To overcome this disadvantage, we utilized a previously well characterized nanoparticle delivery system made of PEG-PGLA polymers [43, 44]. Using this delivery system, we have shown that PP could be encapsulated and delivered to tightly aggregated leukemic spheroids without losing its activity. Also, the PP encapsulated nanoparticles showed specificity in their cytotoxicity by sparing normal mononuclear and stromal cells but efficiently inducing death in ALL cells.

Even though a lot of attention has been given to targeting of glycolysis in cancer cells as a strategy for therapy, a phase I clinical study utilizing a glycolytic inhibitor, 2-deoxy-D-glucose, has been shown to be well tolerated but not very successful in reversing disease progression [45]. Interestingly, inhibitors of the mitochondrial Complex I have now been successfully tested in preclinical studies and are entering clinical trials for use against leukemia and lymphomas (www.clinicaltrials.gov; Identifier: NCT03291938 and NCT02882321). Success of mitochondrial inhibitors has been attributed to increased reliance of tumor cells on both glycolysis and oxidative phosphorylation for its energy needs, and while inhibition of oxidative phosphorylation in normal cells can allow it to switch to glycolysis and still provide for its energy needs, such adaptation is not possible in cancer cells where glycolysis is already upregulated [46]. However, this hypothesis needs to be further tested and confirmed in an in vivo scenario. In the present study, we have demonstrated the capacity of PP to inhibit oxidative phosphorylation accompanied by down-regulation of β-catenin, both of which was sufficient to induce death in the drug resistant PD cell population of ALL. Furthermore, we were successfully able to formulate a nanoparticle drug delivery system that could potentially increase the systemic availability of PP in leukemic patients. In light of our reported data, PP has a promising activity that can be exploited in clinic to treat ALL patients that have relapsed and are refractory to the existing standard-of-care.

## Acknowledgements

The authors would like to acknowledge Kathleen Brundage, director of the WVU Flow Cytometry & Single Cell Core Facility, for her assistance in acquisition of the flow cytometry data and Aniello Infante, lead bioinformatician with the WVU Genomics Core Facility, for his assistance with gene expression analysis. This work was supported by the Alexander B. Osborn Hematopoietic Malignancy and Transplantation Program, Community Foundation for the Ohio Valley Whipkey Trust, NIH grants U54GM104942, P30GM103488, P20GM103434, RO1HL128485, S10OD016165 and P20 GM109098; and AHA grant 17PRE33660333.

